# Objective method reveals minimum time required to quantify net cost of transport across slow to fast walking speeds

**DOI:** 10.1101/2021.09.21.461248

**Authors:** Bolatito Adeyeri, Shernice A. Thomas, Christopher J. Arellano

**Author notes:** <u>Corresponding Author</u>: Christopher J. Arellano, Department of Health and Human Performance, 3875 Holman St., Rm 104 Garrison, University of Houston, Houston, TX 77204-6015, E.

## Abstract

The U-shaped net cost of transport (COT) curve of walking has helped scientists understand the biomechanical basis that underlies energy minimization during walking. However, to produce an individual’s net COT curve, data must be analyzed during periods of steady-rate metabolism. Traditionally, studies analyze the last few minutes of a 6-10 min trial, assuming that steady-rate metabolism has been achieved. Yet, it is possible that an individual achieves steady rates of metabolism much earlier. However, there is no consensus on how to objectively quantify steady-rate metabolism across a range of walking speeds. Therefore, we developed an objective method to determine the minimum time needed for humans to achieve steady rates of metabolism across slow to fast walking speeds. We hypothesized that a shorter time window could be used to produce a net COT curve that is comparable to the net COT curve created using traditional methods. We analyzed metabolic data from twenty-one subjects who completed several 7-min walking trials ranging from 0.50-2.00 m/s. We partitioned the metabolic data for each trial into moving 1-min, 2-min, and 3 min intervals and calculated their slopes. We statistically compared these slope values to values derived from the last 3-min of the 7-min trial, our ‘gold’ standard comparison. We found that a minimum of 2 min is required to achieve steady-rate metabolism and that data from 2-4 min yields a net COT curve that is not statistically different from the one derived from experimental protocols that are generally accepted in the field.

**New and Noteworthy:** A simple slope method reveals that 4-min of walking at speeds between 0.50-1.75 m/s is the minimum time required to produce an accurate net cost of transport curve.

## 1. INTRODUCTION

The U-shaped net cost of transport (COT) curve of walking is a highly conserved feature in humans (Ralston, 1958), which has helped scientists understand the mechanical determinants that underlie metabolic energy minimization during walking (Alexander, 1989; Kuo and Donelan, 2010; Ralston, 1958). Understanding the mechanical determinants that allow humans to minimize their net COT during walking can act as a key measure for diagnosing and treating individuals with gait pathologies (Kuo and Donelan, 2010; Ralston, 1958; Schwartz, 2007; Waters and Mulroy, 1999). However, producing a net COT curve is a time-consuming process that requires each subject to walk across a range of slow to fast walking speeds and reach a steady rate of metabolism at each speed. In past and recent experiments, it has been common for walking trials to last 6-10 minutes with the average metabolic data of the last 2-3 min of the trial being used for analysis (Arellano et al., 2020; Maxwell Donelan et al., 2001; Ralston, 1958). This is done under the assumption that a subject has achieved steady-rate metabolism during this period. However, it is possible that steady-rate metabolism is reached earlier than expected and thus, data collection times could be substantially reduced. Reducing the protocol time for generating a net COT curve can help lessen the overall burden on both experimenter and subject. However, there has been no consensus on how to define steady-rate metabolism to date. Therefore, as a field, we lack a simple, objective method for defining steady-rate metabolism.

For human walking, Duffy et al. (1996) stated that steady-rate metabolism usually occurs after 2 minutes, but it was unclear how they reached this conclusion (Duffy et al., 1996). Since then, there have been several attempts to develop criteria for defining steady-rate metabolism. For example, Schwartz (2007) used Kendall’s tau, a non-parametric rank correlation coefficient, as a statistical means to determine steady rates of oxygen consumption. For reasons unknown however, this approach has not been widely adopted in the field of locomotion energetics. In contrast, other scientists have used a slope method, which quantifies the rate of change in oxygen consumption as a function of time. Plasschaert et al. (2009), for instance, used a slope threshold of 0.00025 ml O_2_·kg^-1^·s^2^ to define steady-rate metabolism. Along similar lines, others (Dennis et al., 2006; Kramer et al., 2018) have defined steady-rate metabolism as a time period when a subject exhibits low variability in their oxygen uptake values across time (a change <10% or ≤2.0 mL O_2_/min). While these variability criteria appear reasonable, there still lacks a methodological explanation as to why these thresholds were selected. This may be the reason why these approaches have not been widely adopted by others. And lastly, others have defined periods of steady-rate metabolism by visual identification of a plateau in the rate of O_2_ consumption (Sims et al., 2018), which fits the classical definition seen in many exercise physiology textbooks (Brooks et al., 2004). In fact, we admit that we have also relied on visually identifying a plateau and suspect that this is a common strategy in the field.

Given this lack of consensus, we developed a systematic approach that would allow us to define steady-rate metabolism and the minimum time required to generate the net COT curve for human walking, which may provide evidence in support of decreasing protocol times. We explored the slope method carried out by Plasschaert et al. (2009) because it is relatively simple and could be easily understood by novice and experienced scientists undertaking measurements of metabolic energy consumption during walking. Based on the prediction of Duffy et al. (1996), we hypothesized that our slope method would identify an earlier time interval that would yield a net COT curve that is not statistically different from the net COT curve produced by the last 3-min of a 7-min walking trial. For practical purposes, we considered a net COT curve produced by the last 3-min of a 7-min walking trial as our ‘gold’ standard, as this reflects a generally accepted methodology in the field of locomotion biomechanics and energetics.

## 2. MATERIALS AND METHODS

### Participants and Experimental protocol

Twenty-one young, healthy adults (13 W, 8M) participated in this study (age = 25.38±2.92 years, mass = 68.37±12.41 kg, height = 1.70±0.09 m) (Thomas et al., 2021). They were non-smokers and physically active per ACSM guidelines (American College of Sports Medicine Guidelines, 2018), with a body mass index <30.0. All participants gave written informed consent as per the University of Houston Institutional Review Board. The experiment began by measuring subjects’ standing metabolic energy consumption for 7 minutes using a metabolic cart (ParvoMedics TrueMax2400, UT, USA) that reported the average expired oxygen consumption (V□O_2_) and carbon dioxide (V□CO_2_) every 15 seconds.

The experiment began by measuring subjects’ standing metabolic energy consumption for 7 minutes using a metabolic cart (Parvo Medics TrueMax2400, UT, USA), with a setting that reported average rates of oxygen consumption (V□O_2_) and carbon dioxide (V□CO_2_) approximately every 15 seconds. In brief, the average data are calculated from the accumulated *V*O_2_ which starts with the beginning of the first detectable breath and ends when a complete breath that is detected close to or after it exceeds 15 seconds. Therefore, the average data is quantified by dividing the accumulated VO_2_ by the accumulated time window of roughly 15 seconds (personal communication, Pat Yeh, Ph.D., Chief Architect, Parvo Medics). The same procedure is followed for average V□CO_2_. We followed the guidelines for metabolic testing described by the Parvo Medics manual, starting with a 3-L syringe flow calibration procedure. The flow calibration consisted first of defining room temperature, barometric pressure, and relative humidity. Then, the 3-L syringe was used to achieve slow to fast peak stroke rates, ranging between 50-80, 100-199, 200-299, 300-399, and 400-499 L/m. We continued the flow calibration until reaching a ± 0.1% difference between the average volume measured during the calibration and the known volume of the 3-L syringe. This was usually achieved within two attempts. After successful flowmeter calibration, gas analyzers were calibrated with a gas tank with known percentages of CO_2_ and O_2_. Again, this process was continued until the gas concentration readings from the metabolic cart matched the known gas tank concentrations. This was also achieved within two attempts.

The subjects walked on a dual-belt treadmill (Bertec Corp, OH, USA), completing the 7-minute trials in a randomized order for all seven speeds, which ranged from 0.50 to 2.00 m/s in 0.25 m/s intervals (Figure 1). In order to minimize the effects of fatigue, participants rested for 5 minutes between the trials. The respiratory exchange ratio (RER) was monitored for each subject to ensure values remained <1.00, indicating that metabolic energy was provided primarily by aerobic pathways. If any subject was unable to keep up with the treadmill or produced an RER above 1.00, the trial was stopped and excluded from data analysis. For instance, at 1.00 m/s, one subject did not reach a steady rate of metabolism. In addition, at 1.75 m/s and 2.00 m/s, two and nine subjects could not keep up with the treadmill, respectively.

**Figure 1.**
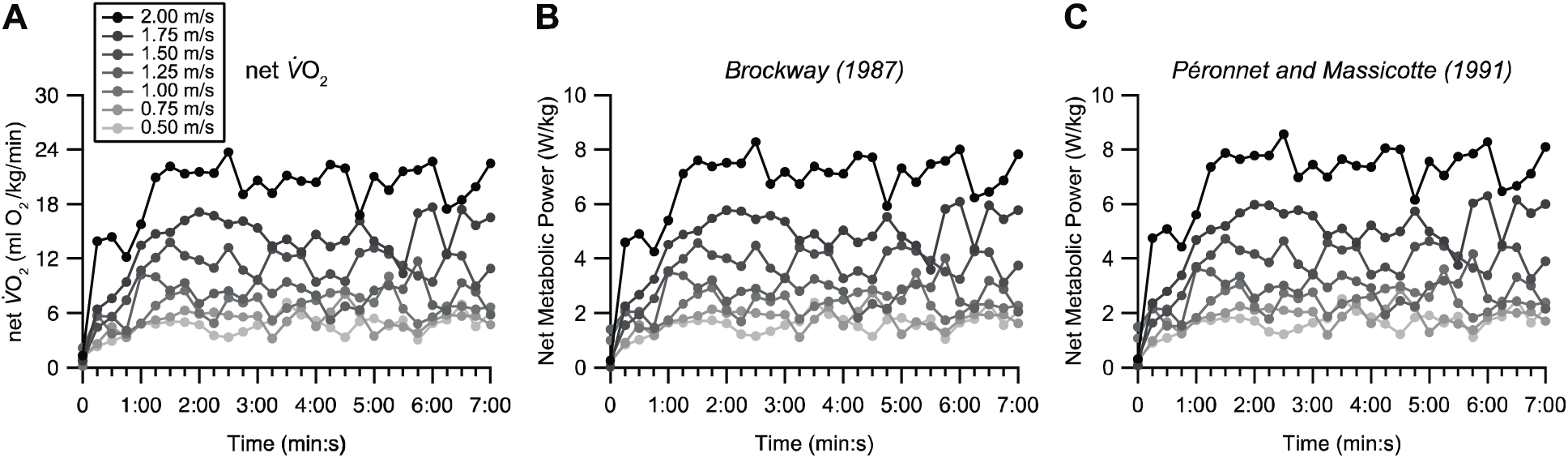
Representative time-series data from a single subject illustrates the demand for net oxygen consumption (A) and net metabolic power (B and C) while walking across slow to fast speeds. The first data points at zero marks the value for the 15 seconds before the start of the seven-minute trial. Note the sharp rise in the demand for metabolic energy when the subject starts to walk at moderate and fast speeds (≥1.25 m/s). Even though the energy demand increases steadily, the subject reaches a steady rate of metabolism at 2-min.

## Data Analysis

We used the average V□O_2_ and V□CO_2_ values to calculate metabolic power for each standing and walking trial using the Brockway equation (Brockway, 1987) and the Péronnet and Massicotte equation (Kipp et al., 2018; Péronnet and Massicotte, 1991). For each speed, we calculated net metabolic power by subtracting each participant’s standing metabolic power from their gross metabolic power values in each walking trial (Figure 1). To mirror typical data collection methods, we divided the seven-minute trials into overlapping 3-minute, 2-minute, and 1-minute intervals. For the 3-minute analyses, for example, the first window started at 15 sec and ended at 180 sec. Note that the 15 sec mark represents the average metabolic data sampled between the start of the walking trial and the subsequent 15 sec. For simplicity we refer to the first window as 0-3:00 min:s. Then, this window moved to the next time point, starting at 30 sec and ending at 195 sec. The 30 sec mark represents the average metabolic data sampled between the end of the first 15 sec of the trial and the following 15 sec. We refer to the second window as 0:15-3:15 min:s. This process continued until reaching the end of the entire 7-minute time-series. This analysis was replicated for the 2-minute and 1-minute intervals. This resulted in seventeen intervals lasting 3 minutes each, twenty-one intervals lasting 2 minutes each, and twenty-five intervals lasting 1 minute each. For each subject and speed, we used the time-series intervals to quantify the slope ([ml O_2_/kg/min]/s or [W/kg]/s) using linear regression. For subsequent analyses, we also quantified the net cost of transport by dividing net V□O_2_ (ml O_2_/kg/min *x* 1 min/60 s) by walking speed (m/s) and net metabolic power (W/kg) by walking speed (m/s). We used MATLAB (Mathworks, Inc., MA, USA) to perform all computational calculations and descriptive analyses.

## Statistical Analysis

For each 1-min, 2-min, and 3-min interval, we aggregated the data at each speed for all subjects and then calculated the regression line that estimated the change in slope as a function of speed (i.e., 0.5 m/s to 2.0 m/s; see Figure 2). For instance, a regression line was derived for each 1-min interval, starting from 0-1:00 min:s, then from 0:15 to 1:15 min:s, then from 0:30 to 1:30 min:s and so on, until reaching the final 6:00-7:00 min:s. The same approach was carried out for the 2-min and 3-min intervals. We then used the Dunnett method of multiple comparison (MC) to determine whether each regression line was statistically different from the regression line that was based on the last 3-min of the 7-min trial, predesignated here as the control. The criteria for significance were based on a directional one-tailed test, an alpha value equal to 0.05, *k* number of comparisons, and *v* degrees of freedom based on the residual sum of squares. To calculate the critical *t*-value for the Dunnett multiple comparison method, we used the ‘nCDunnett’ package provided in R-software, which yielded the following results: *t*_1-min_ = 2.721 (*k* = 25, *v* = 266), *t*_2-min_ = 2.674 (*k* = 21, *v* = 266), and *t*_3-min_ = 2.598 (*k* = 16, *v* = 266).

**Figure 2.**
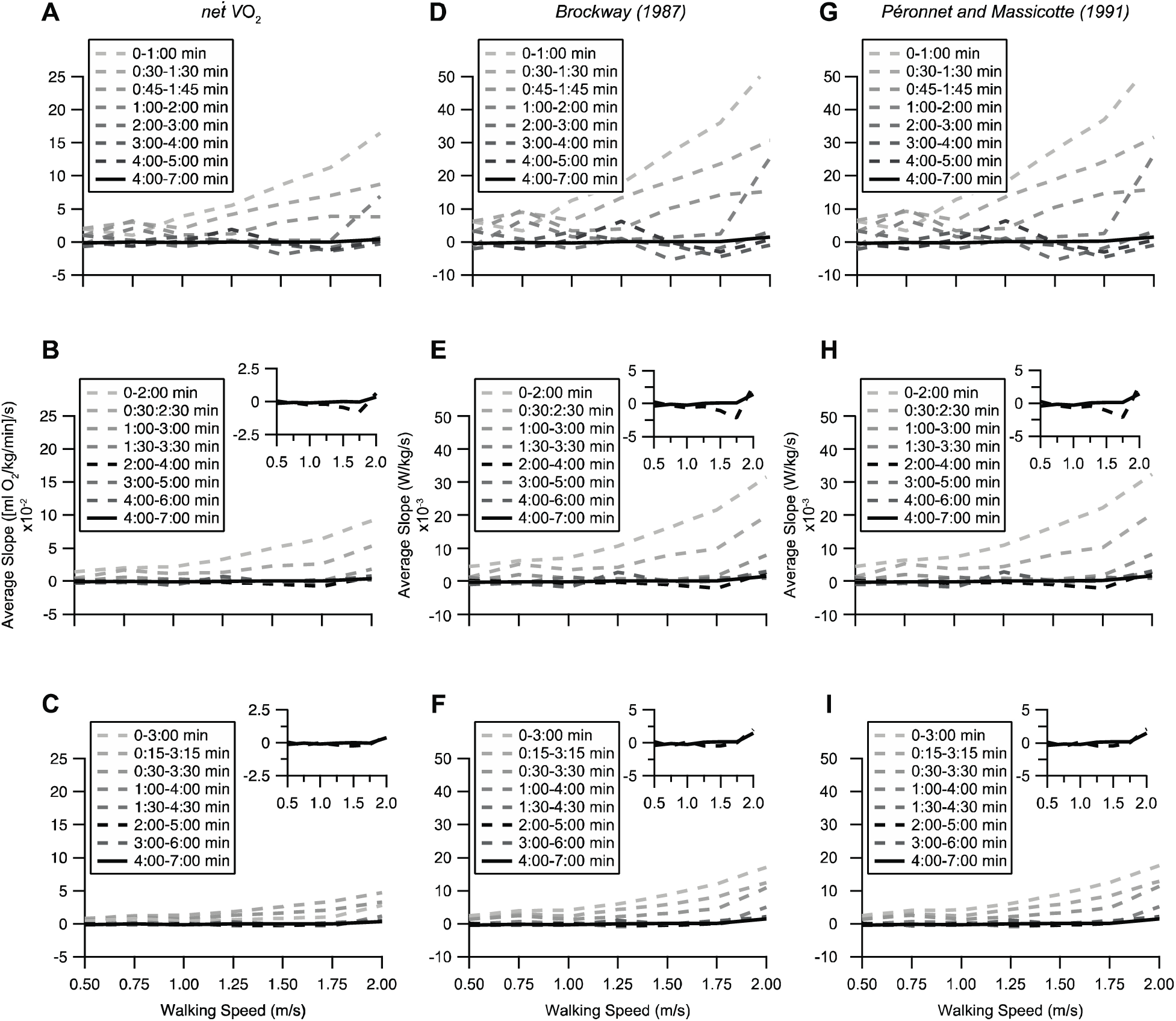
The trends illustrate the changes in the average slope quantified for a 1-min (A, D, G), 2-min (B, E, H), and 3-min (C, F, I) time window. The slope at each walking speed represents the change in net oxygen consumption or net metabolic power over the specific time window across the time-series data. By plotting the slope as a function of speed (using the individual data points), a regression line was fit across all the 1-min intervals, starting from 0-1:00 min:s, then 0:15 to 1:15 min:s, then 0:30 to 1:30 min:s and so on, until reaching the final 6:00-7:00 min:s. The same procedure was followed for the 2-min and 3-min intervals. The trends show that as the time window moved along, toward the end of the original time-series data, the unit increase in slope decreased, eventually reaching similar values to those observed from the 4-7 min time window, our ‘gold’ standard (solid black line). To allow for closer inspection, the insets for the 2-min and 3-min graphs highlight the earliest time window that is not statistically different from the ‘gold’ standard. No inset is included for the 1-min intervals as they were not considered for curve comparisons. For clarity, several time intervals were omitted; however, those omitted exhibited the same downward trend.

Based on the outcomes of the Dunnett method, we moved forward with a non-linear regression analysis using R software to compare the rate of oxygen consumption, net metabolic power, and net COT curves derived from a 2-4 min and 2-5 min window to that derived from the last 4-7 min window (see Discussion for details). We then followed up with pre-planned comparisons between the average net COT values at each level of speed using paired *t*-tests with α = 0.05 (SPSS Inc., Chicago, IL). Note that for each level of speed, where values were compared using paired t-test, sample sizes were always equal. If the data did not meet the assumptions of normality, we used the non-parametric Wilcoxon signed-rank test. For clarity, the average and standard deviation values for net COT at each speed and time window are provided in Table 1A-C while the regression equations signifying the slope, intercept, adjusted coefficient of determination (*r*^2^ _adj_), along with the *F* and *p* values are provided in Table 2.

**Table 1A.** Net COT values derived from net *V*□O_2_ data extracted from the selected time windows at each speed.

| Walking Speed<br>(m/s) | Net COT (ml $O_2$/kg/m)<br>for 2-min interval<br>(2.00-4.00 min) | Net COT (ml $O_2$/kg/m)<br>for 3-min interval<br>(2.00-5.00 min) | Net COT (ml $O_2$/kg/m)<br>for 'gold' standard<br>(4.00-7.00 min) |
| --- | --- | --- | --- |
| 0.50 | 0.132 ± 0.026 ($n=21$)<br>$d = 0.306$<br>$p = 0.322$ (WS) | 0.132 ± 0.027 ($n=21$)<br>$d = 0.479$<br>$p = 0.040$ | 0.129 ± 0.026 ($n=21$) |
| 0.75 | 0.113 ± 0.022 ($n=21$)<br>$d = 0.406$<br>$p = 0.078$ | 0.112 ± 0.022 ($n=21$)<br>$d = 0.414$<br>$p = 0.050$ (WS) | 0.109 ± 0.020 ($n=21$) |
| 1.00 | 0.103 ± 0.017 ($n=20$)<br>$d = 0.108$<br>$p = 0.634$ | 0.104 ± 0.016 ($n=20$)<br>$d = 0.274$<br>$p = 0.236$ | 0.102 ± 0.016 ($n=20$) |
| 1.25 | 0.105 ± 0.017 ($n=21$)<br>$d = 0.327$<br>$p = 0.150$ | 0.104 ± 0.016 ($n=21$)<br>$d = 0.259$<br>$p = 0.250$ | 0.103 ± 0.014 ($n=21$) |
| 1.50 | 0.115 ± 0.016 ($n=21$)<br>$d = 0.423$<br>$p = 0.062$ (WS) | 0.114 ± 0.016 ($n=21$)<br>$d = 0.353$<br>$p = 0.085$ (WS) | 0.113 ± 0.015 ($n=21$) |
| 1.75 | 0.131 ± 0.020 ($n=19$)<br>$d = -0.204$<br>$p = 0.385$ | 0.132 ± 0.195 ($n=19$)<br>$d = -0.201$<br>$p = 0.494$ (WS) | 0.132 ± 0.020 ($n=19$) |
| 2.00 | 0.155 ± 0.018 ($n=12$)<br>$d = -0.942$<br>**$p = 0.008$** | 0.156 ± 0.018 ($n=12$)<br>$d = -1.242$<br>**$p = 0.001$** | 0.161 ± 0.017 ($n=12$) |

All comparisons at each speed were made between an earlier interval and the 'gold' standard interval with Bonferroni adjusted significance set at $P < 0.025$ . WS indicates $p$ value from Wilcoxon Signed Rank Test and bold $p$ value denotes significant differences when compared to the 'gold' standard.
Values expressed as mean ± SD
Effect size calculated based on Cohen's $d = t/\sqrt{n}$

**Table 2.** Best fit curve equations and mean square error of the regression fits for the time windows displayed in Figure 4.

|  | Net VO<sub>2</sub> | Brockway (1987) | Péronnet & Massicotte (1991) |
| --- | --- | --- | --- |
| 2-4 min | Net VO<sub>2</sub> (ml O<sub>2</sub>/kg/min) =<br>(5.507 ± 0.597)speed<sup>2</sup> –<br>(5.426 ± 1.470)speed + (5.507 ± 0.813), $R^2_{adj} = 0.904$,<br>$F_{(2,132)} = 632.46$, $p < 0.001$.<br>MSE<sub>2-4min</sub> = 2.050 | Net Metabolic Power (W/kg) =<br>(2.138 ± 0.205)speed<sup>2</sup> –<br>(2.118 ± .505)speed + (1.967 ± 0.279), $R^2_{adj} = 0.906$,<br>$F_{(2,132)} = 650.10$, $p < 0.001$.<br>MSE<sub>2-4min</sub> = 0.242 | Net Metabolic Power (W/kg) =<br>(2.205 ± 0.212)speed<sup>2</sup> –<br>(2.190 ± 0.521)speed + (2.063 ± 0.288), $R^2_{adj} = 0.906$,<br>$F_{(2,132)} = 649.07$, $p < 0.001$.<br>MSE<sub>2-4min</sub> = 0.257 |
| 2-5 min | Net VO<sub>2</sub> (ml O<sub>2</sub>/kg/min) =<br>(6.058 ± 0.586)speed<sup>2</sup> –<br>(5.798 ± 1.442)speed + (5.653 ± 0.797), $R^2_{adj} = 0.909$,<br>$F_{(2,132)} = 669.067$, $p < 0.001$.<br>MSE<sub>2-5min</sub> = 1.973 | Net Metabolic Power (W/kg) =<br>(2.203 ± 0.202)speed<sup>2</sup> –<br>(2.248 ± 0.498)speed + (2.020 ± 0.275), $R^2_{adj} = 0.911$,<br>$F_{(2,132)} = 685.30$, $p < 0.001$.<br>MSE<sub>2-5min</sub> = 0.235 | Net Metabolic Power (W/kg) =<br>(2.273 ± 0.208)speed<sup>2</sup> –<br>(2.324 ± 0.513)speed + (2.117 ± 0.284), $R^2_{adj} = 0.911$,<br>$F_{(2,132)} = 683.91$, $p < 0.001$.<br>MSE<sub>2-5min</sub> = 0.250 |
| 4-7 min | Net VO<sub>2</sub> (ml O<sub>2</sub>/kg/min) =<br>(6.584 ± 0.571)speed<sup>2</sup> –<br>(6.812 ± 1.408)speed + (5.983 ± 0.778), $R^2_{adj} = 0.918$,<br>$F_{(2,132)} = 751.320$, $p < 0.001$.<br>MSE<sub>4-7min</sub> = 1.879 | Net Metabolic Power (W/kg) =<br>(2.378 ± 0.198)speed<sup>2</sup> –<br>(2.564 ± 0.488)speed + (2.122 ± 0.270), $R^2_{adj} = 0.920$,<br>$F_{(2,132)} = 768.75$, $p < 0.001$.<br>MSE<sub>4-7min</sub> = 0.226 | Net Metabolic Power (W/kg) =<br>(2.452 ± 0.204)speed<sup>2</sup> –<br>(2.646 ± 0.504)speed + (2.221 ± 0.278), $R^2_{adj} = 0.920$,<br>$F_{(2,132)} = 767.35$, $p < 0.001$.<br>MSE<sub>4-7min</sub> = 0.240 |
| 2-4 min | Net COT (ml O<sub>2</sub>/kg/m) =<br>(0.069 ± 0.008)speed<sup>2</sup> –<br>(0.157 ± 0.020)speed + (0.192 ± 0.011), $R^2_{adj} = 0.374$,<br>$F_{(2,132)} = 41.007$, $p < 0.001$.<br>MSE<sub>2-4min</sub> = 0.000388 | Net COT (J/kg/m) =<br>(1.476 ± 0.166)speed<sup>2</sup> –<br>(3.292 ± 0.408)speed + (3.925 ± 0.225), $R^2_{adj} = 0.412$,<br>$F_{(2,132)} = 47.88$, $p < 0.001$.<br>MSE<sub>2-4min</sub> = 0.158 | Net COT (J/kg/m) =<br>(1.551 ± 0.171)speed<sup>2</sup> –<br>(3.496 ± 0.420)speed + (4.151 ± 0.232), $R^2_{adj} = 0.411$,<br>$F_{(2,132)} = 47.81$, $p < 0.001$.<br>MSE<sub>2-4min</sub> = 0.167 |
| 2-5 min | Net COT (ml O<sub>2</sub>/kg/m) =<br>(0.071 ± 0.008)speed<sup>2</sup> –<br>(0.160 ± 0.020)speed + (0.193 ± 0.011), $R^2_{adj} = 0.396$,<br>$F_{(2,132)} = 44.935$, $p < 0.001$.<br>MSE<sub>2-5min</sub> = 0.000376 | Net COT (J/kg/m) =<br>(1.514 ± 0.164)speed<sup>2</sup> –<br>(3.367 ± 0.403)speed + (3.953 ± 0.223), $R^2_{adj} = 0.433$,<br>$F_{(2,132)} = 52.14$, $p < 0.001$.<br>MSE<sub>2-5min</sub> = 0.154 | Net COT (J/kg/m) =<br>(1.591 ± 0.169)speed<sup>2</sup> –<br>(3.574 ± 0.415)speed + (4.180 ± 0.229), $R^2_{adj} = 0.432$,<br>$F_{(2,132)} = 52.06$, $p < 0.001$.<br>MSE<sub>2-5min</sub> = 0.163 |
| 4-7 min | Net COT (ml O<sub>2</sub>/kg/m) =<br>(0.074 ± 0.008)speed<sup>2</sup> –<br>(0.164 ± 0.019)speed + (0.192 ± 0.011), $R^2_{adj} = 0.459$,<br>$F_{(2,132)} = 57.905$, $p < 0.001$.<br>MSE<sub>4-7min</sub> = 0.000342 | Net COT (J/kg/m) =<br>(1.573 ± 0.158)speed<sup>2</sup> –<br>(3.427 ± 0.389)speed + (3.924 ± 0.215), $R^2_{adj} = 0.491$,<br>$F_{(2,132)} = 65.64$, $p < 0.001$.<br>MSE<sub>4-7min</sub> = 0.143 | Net COT (J/kg/m) =<br>(1.650 ± 0.163)speed<sup>2</sup> –<br>(3.635 ± 0.401)speed + (4.151 ± 0.222), $R^2_{adj} = 0.488$,<br>$F_{(2,132)} = 64.93$, $p < 0.001$.<br>MSE<sub>4-7min</sub> = 0.153 |

**Table 2B.** Net COT values derived from net metabolic power data extracted from the selected time windows at each speed. Net metabolic power is estimated from the equation published by Brockway (1987).

| Walking Speed<br>(m/s) | Net COT (J/kg/m)<br>for 2-min interval<br>(2.00-4.00 min) | Net COT (J/kg/m)<br>for 3-min interval<br>(2.00-5.00 min) | Net COT (J/kg/m)<br>for 'gold' standard<br>(4.00-7.00 min) |
| --- | --- | --- | --- |
| 0.50 | 2.663 ± 0.531 (*n*=21)<br>*d* = 0.233<br>*p* = 0.298 | 2.666 ± 0.543 (*n*=21)<br>*d* = 0.395<br>*p* = 0.085 | 2.617 ± 0.537 (*n*=21) |
| 0.75 | 2.271 ± 0.437 (*n*=21)<br>*d* = 0.359<br>*p* = 0.115 | 2.251 ± 0.429 (*n*=21)<br>*d* = 0.380<br>*p* = 0.097 | 2.210 ± 0.393 (*n*=21) |
| 1.00 | 2.087 ± 0.338 (*n*=20)<br>*d* = 0.038<br>*p* = 0.865 | 2.102 ± 0.318 (*n*=20)<br>*d* = 0.190<br>*p* = 0.406 | 2.081 ± 0.317 (*n*=20) |
| 1.25 | 2.125 ± 0.327 (*n*=21)<br>*d* = 0.149<br>*p* = 0.502 | 2.106 ± 0.307 (*n*=21)<br>*d* = -0.026<br>*p* = 0.906 | 2.108 ± 0.290 (*n*=21) |
| 1.50 | 2.325 ± 0.328 (*n*=21)<br>*d* = 0.069<br>*p* = 0.614 (WS) | 2.322 ± 0.332 (*n*=21)<br>*d* = 0.028<br>*p* = 0.821 (WS) | 2.320 ± 0.30 (*n*=21) |
| 1.75 | 2.692 ± 0.417 (*n*=19)<br>*d* = -0.342<br>*p* = 0.153 | 2.711 ± 0.406 (*n*=19)<br>*d* = -0.380<br>*p* = 0.115 | 2.743 ± 0.418 (*n*=19) |
| 2.00 | 3.226 ± 0.397 (*n*=12)<br>*d* = -1.116<br>***p* = 0.003** | 3.253 ± 0.395 (*n*=12)<br>*d* = -1.388<br>***p* = 0.001** | 3.359 ± 0.372 (*n*=12) |

All comparisons at each speed were made between an earlier interval and the 'gold' standard interval with Bonferroni adjusted significance set at $P < 0.025$ . WS indicates *p* value from Wilcoxon Signed Rank Test and bold *p* value denotes significant differences when compared to the 'gold' standard.
Values expressed as mean ± SD
Effect size calculated based on Cohen's $d = t/\sqrt{n}$

**Table 3C.** Net COT values derived from net metabolic power data extracted from the selected time windows at each speed. Net metabolic power is estimated from the equation published by Péronnet & Massicotte (1991).

| Walking Speed<br>(m/s) | Net COT (J/kg/m)<br>for 2-min interval<br>(2.00-4.00 min) | Net COT (J/kg/m)<br>for 3-min interval<br>(2.00-5.00 min) | Net COT (J/kg/m)<br>for 'gold' standard<br>(4.00-7.00 min) |
| --- | --- | --- | --- |
| 0.50 | 2.809 ± 0.546 (*n*=21)<br>*d* = 0.229<br>*p* = 0.306 | 2.81 ± 0.56 (*n*=21)<br>*d* = 0.389<br>*p* = .090 | 2.763 ± 0.553 (*n*=21) |
| 0.75 | 2.382 ± 0.451 (*n*=21)<br>*d* = 0.357<br>*p* = 0.117 | 2.362 ± 0.442 (*n*=21)<br>*d* = 0.377<br>*p* = 0.099 | 2.320 ± 0.405 (*n*=21) |
| 1.00 | 2.182 ± 0.348 (*n*=20)<br>*d* = 0.035<br>*p* = 0.877 | 2.197 ± 0.328 (*n*=20)<br>*d* = 0.186<br>*p* = 0.416 | 2.176 ± 0.327 (*n*=20) |
| 1.25 | 2.213 ± 0.336 (*n*=21)<br>*d* = 0.141<br>*p* = 0.524 | 2.194 ± 0.315 (*n*=21)<br>*d* = -0.038<br>*p* = 0.862 | 2.197 ± 0.298 (*n*=21) |
| 1.50 | 2.415 ± 0.338 (*n*=21)<br>*d* = 0.053<br>*p* = 0.689 (WS) | 2.412 ± 0.342 (*n*=21)<br>*d* = 0.013<br>*p* = 0.876 (WS) | 2.411 ± 0.311 (*n*=21) |
| 1.75 | 2.791 ± 0.430 (*n*=19)<br>*d* = -0.348<br>*p* = 0.147 | 2.812 ± 0.418 (*n*=19)<br>*d* = -0.388<br>*p* = 0.108 | 2.844 ± 0.431 (*n*=19) |
| 2.00 | 3.339 ± 0.410 (*n*=12)<br>*d* = -1.123<br>***p* = 0.003** | 3.367 ± 0.409 (*n*=12)<br>*d* = -1.392<br>***p* = 0.001** | 3.477 ± 0.385 (*n*=12) |

All comparisons at each speed were made between an earlier interval and the 'gold' standard interval with Bonferroni adjusted significance set at $P < 0.025$ . WS indicates *p* value from Wilcoxon Signed Rank Test and bold *p* value denotes significant differences when compared to the 'gold' standard.
Values expressed as mean ± SD
Effect size calculated based on Cohen's $d = t/\sqrt{n}$

## 3. RESULTS

### Influence of Time Window across Speed

The regression lines based on the initial time window of 1-min, 2-min, and 3-min intervals revealed that the slope significantly increased as a function of speed (Fig. 2). For example, the slope derived for a 1-min interval was closer to zero for the slowest speed (0.5 m/s) and increased to its highest value at 2.0 m/s. As the time window moved along, toward the end of the time-series data, the unit increase in the magnitude of the slope decreased, and therefore, the linear relationship between slope and speed became less steep. In general, the linear relationship between the magnitude of the slope and speed for the last 2-min and 1-min time windows overlapped closely with the last 3-min window, i.e., the control.

#### Change in slope based on Net V□O_2_

The regression lines for the initial and last time window for the 1-min intervals were slope = 0.094(speed) – 0.049, (*r*^2^ _adj_) = 0.143 and slope = 0.010(speed) – 0.010, (*r*^2^ _adj_) = 0.012, respectively. The regression lines for the initial and last time window for the 2-min intervals were slope = 0.047(speed) – 0.018, (*r*^2^ _adj_) = 0.412 and slope = 0.002(speed) – 0.003, (*r*^2^ _adj_) = -0.002, respectively. And finally, the regression lines for the initial and last time window for the 3-min intervals were slope = 0.024(speed) – 0.007, (*r*^2^ _adj_) = 0.369 and slope = 0.002(speed) – 0.003, (*r*^2^ _adj_) = 0.017, respectively.

#### Change in slope based on Net Metabolic Power via Brockway (1987)

The regression lines for the initial and last time window for the 1-min intervals were slope = 0.030(speed) – 0.016, (*r*^2^ _adj_) = 0.416 and slope = 0.003(speed) - 0.003, (*r*^2^ _adj_) = 0.013, respectively. The regression lines for the initial and last time window for the 2-min intervals were slope = 0.016(speed) – 0.007, (*r*^2^ _adj_) = 0.442 and slope = 0.001(speed) - 0.001, (*r*^2^ _adj_) 0.001, respectively. And finally, the regression lines for the initial and last time window for the 3-min intervals were slope = 0.009(speed) – 0.003, (*r*^2^ _adj_) = 0.429 and slope = 0.001(speed) - 0.001, (*r*^2^ _adj_) = 0.027, respectively.

#### Change in slope based on Net Metabolic Power via from Péronnet & Massicotte (1991)

To illustrate using the Péronnet & Massicotte data, the regression lines for the initial and last time window for the 1-min intervals were slope = 0.031(speed) – 0.016, (*r*^2^ _adj_) = 0.416 and slope = 0.004(speed) – 0.003, (*r*^2^ _adj_) = 0.013, respectively. The regression lines for the initial and last time window for the 2-min intervals were slope = 0.017(speed) – 0.007, (*r*^2^ _adj_) = 0.444 and slope = 0.001(speed) - 0.001, (*r*^2^ _adj_) = 0.001, respectively. And finally, the regression lines for the initial and last time window for the 3-min intervals were slope = 0.009(speed) – 0.003, (*r*^2^ _adj_) = 0.431 and slope = 0.001(speed) – 0.001, (*r*^2^ _adj_) = 0.028, respectively.

### Dunnett’s MC Method to Determine Earliest Time Window

As shown in Fig. 3, the comparisons between the regression lines revealed that for 1-min window comparisons, the *t*-statistic fell below the critical *t*-value at an interval between 1:15-2:15 min; however, when contrasted with the 2-min and 3-min windows, the *t*-statistic exhibited greater fluctuations for the subsequent comparisons against the control. For a 2-min window, comparisons of regression lines revealed that a window between 1:30-3:30 min was not statistically different from the last 3-min window. And finally, for a 3-min window, comparison of regression lines revelated that a window between 2:00-5:00 min was not significantly different from the last 3-min. Given these observations, we chose to compare the 3-min window defined from 2:00-5:00 min against the control. And to keep consistent with our 3-min window comparison, we also chose to compare the 2-min window defined from 2:00-4:00 min against the control. Although the *t*-statistic fell below the Dunnett’s critical t-value for the 1-min window, we avoided any regression fit comparisons using this window due to higher fluctuations in the pattern.

**Figure 3.**
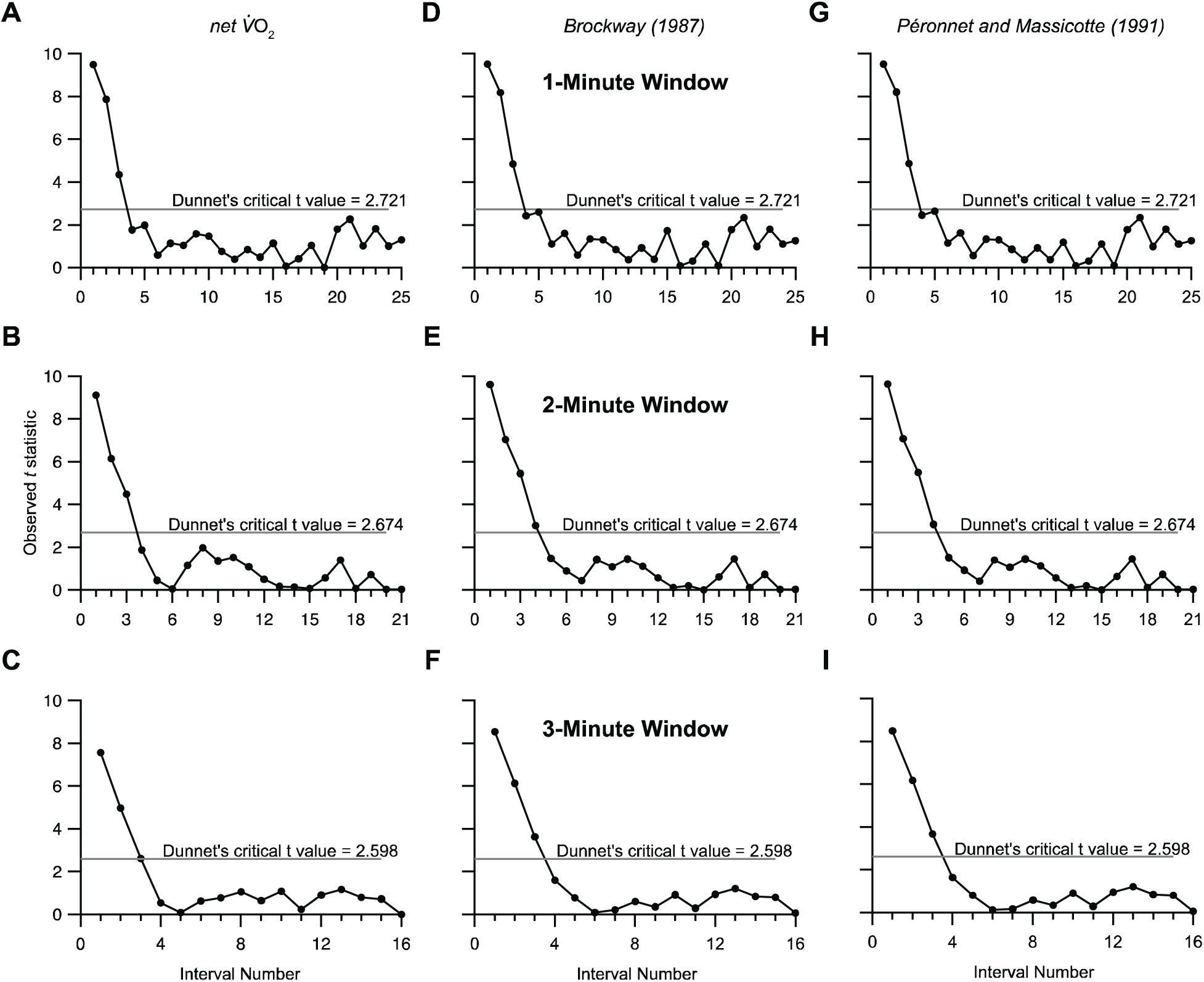
The trends illustrate changes in the observed *t*-statistic derived from systematic comparisons between regression lines, where each interval, from start to end, is compared against the control, defined as the last 3-min of a 7-min walking trial. In general, the observed *t*-statistic is highest for the first interval, then trends downward until reaching a value that is below Dunnett’s critical *t*-statistic, indicating that subjects have achieved a steady rate of metabolism that is not statistically different from the last 3-min of the 7-min trial. Interval number 1 represents the time window 0:00-1:00 min:s and interval number 2 represent the 0:15-1:15 min:s, and so on. Regression line comparisons revealed that the 1-min window between 0:45-1:45 min:s (interval number 4) reached a value below threshold (A, D, G), but the observed *t*-statistic had higher fluctuations in the pattern. In contrast, regression line comparisons revealed that a 2-min window between 1:00-3:00 min:s (interval number 5) (B, E, H) and a 3-min window between 0:45-3:45 min:s (interval number 4) (C, F, I) reached a value below Dunnett’s threshold, and remained consistent for subsequent comparisons. Overall, the observed *t*-statistic trends for 2-min and 3-min windows demonstrate that subjects reach a steady rate of metabolism much earlier that the last 3-min of a 7-min walking trial.

### Net V□O2, Net Metabolic Power, and Net Cost of Transport Curves

The nonlinear regression line comparisons characterizing the net V□O_2_ as a function of speed did not significantly differ between the last 3-min interval and the 3-min interval between 2:00-5:00 min or between the last 3-min interval and the 2-min interval between 2:00-4:00 min (Figure 4; *p* values provided in the caption). In addition, the nonlinear regression line comparisons characterizing the net COT as a function of speed did not significantly differ between the last 3-min interval and the 3-min interval between 2:00-5:00 min or between the last 3-min interval and the 2-min interval between 2:00-4:00 min. The same results were observed when carrying out the nonlinear comparisons for net metabolic power and net COT, regardless of whether these data were derived using the Brockway equation or the Péronnet & Massicotte equation (Figure 5 and 6; *p* values provided in the captions). Follow-up pairwise comparisons between the last 3-min interval and the earlier time intervals showed that the net COT values were not statistically different at speeds of 0.5 m/s to 1.75 m/s (all *p*’s > 0.05). However, when compared to the last 3-min interval, the net COT values for the fastest speed of 2.00 m/s were consistently lower for the earlier time intervals (Table 1).

**Figure 4.**
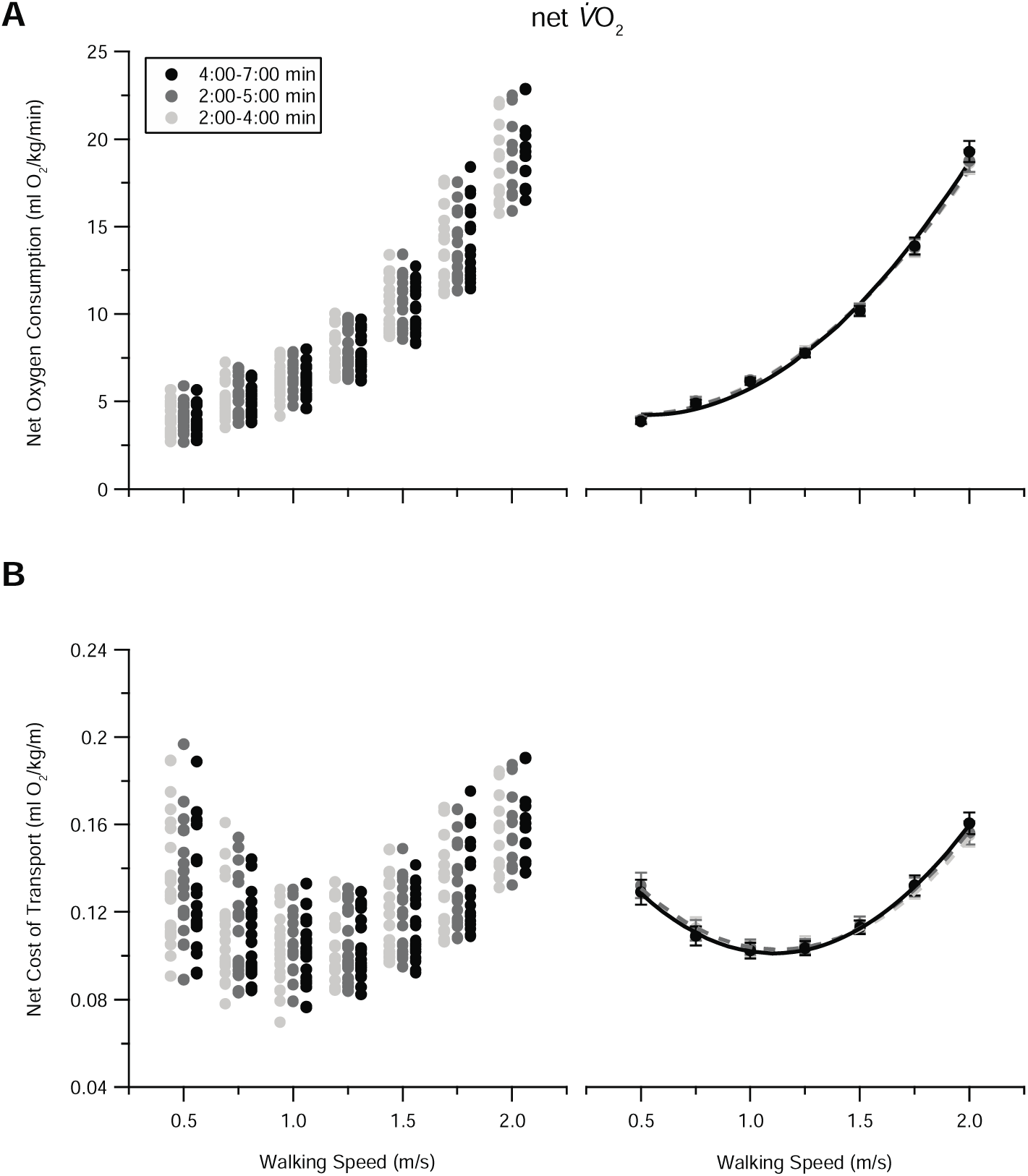
(A) The demand for net oxygen consumption and (B) net cost of transport changes nonlinearly with walking speed, with estimates based on the average rates of oxygen consumption measured by Parvo Medics TrueMax2400 system. The left column represents the original data points, and the right column represents the regression curve fits to these data. The data in the right column represent the mean ± sem. As expected, the demand for net oxygen consumption increases curvilinearly with speed. In contrast, the net cost of transport exhibits a U-shaped curve, highlighting the observation that walking at a speed between 1-1.3 m/s minimizes the net cost of transport, i.e., the net oxygen cost required to move one kilogram of body mass a unit meter. For ease of visual inspection, a small offset is applied to the original data points along the abscissa in the left column of (A) and (B). Statistical comparisons made from the regression equations characterizing the non-linear relationship between net oxygen consumption vs. speed and net cost of transport vs. speed were not statistically different for the time windows taken from 2-4 min, 2-5 min, and 4-7 min (net oxygen consumption comparisons: *p*_2-4min vs 4-7min_ = 0.741 and *p*_2-5min vs 4-7min_ = 0.796; net COT comparisons: *p*_2-4min vs 4-7min_ = 0.974 and *p*_2-5min vs 4-7min_ = 0.972). Note that the regression curves for the 2-4 min window overlap with the 2-5 min window, therefore, they are indistinguishable from one another. The equations characterizing the polynomial non-linear regression fits in the right column are listed in Table 2.

**Figure 5.**
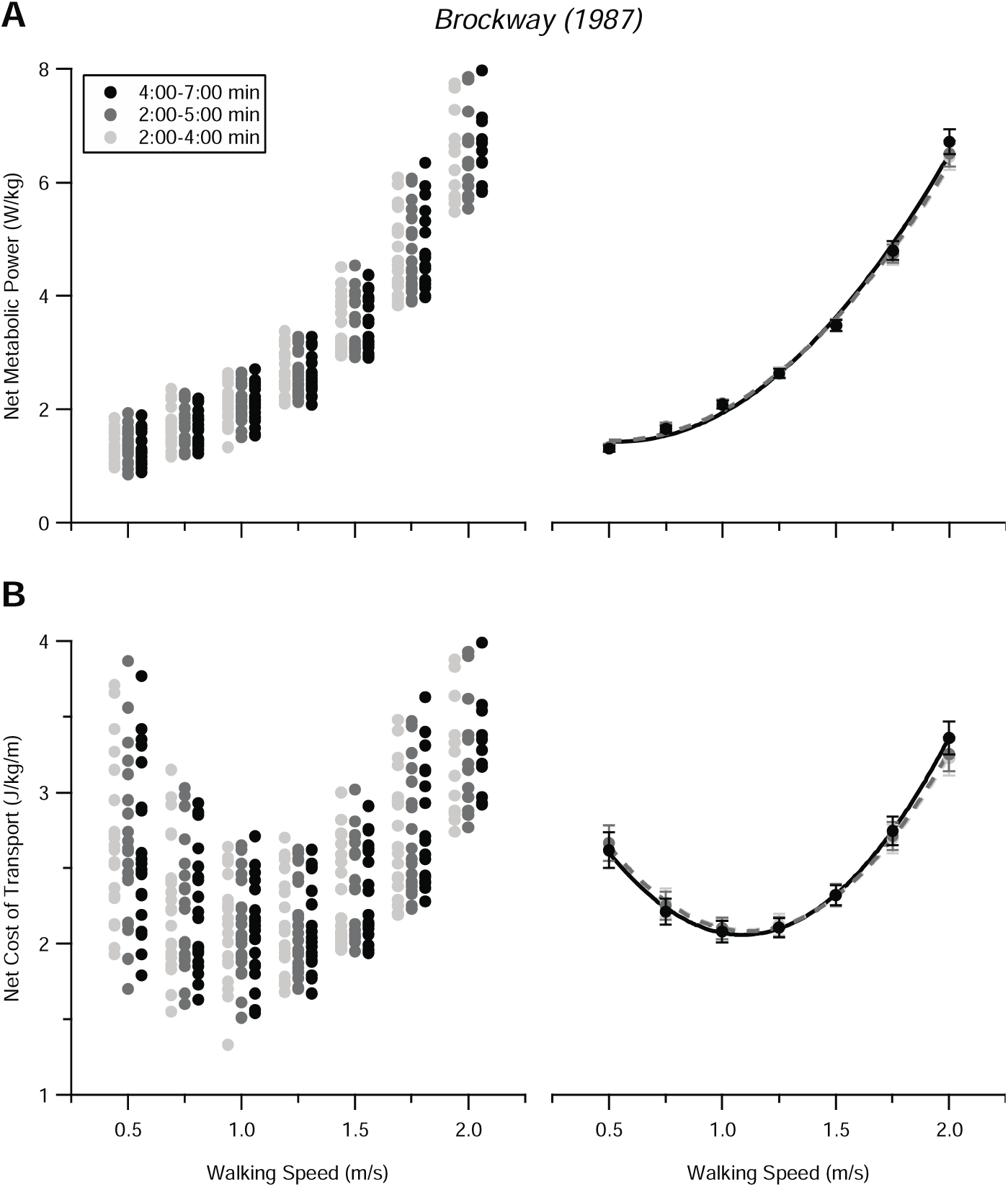
(A) The demand for net metabolic power and (B) net cost of transport changes nonlinearly with walking speed. The left column represents the original data points that are estimated from the Brockway (1987) equation, and the right column represents the regression curves fit to these data. The data in the right column represent the mean ± sem. Similar to the trends and interpretation in Figure 4, the demand for net metabolic power increases curvilinearly with speed while the net cost of transport exhibits a U-shaped curve, highlighting the observation that walking at a speed between 1-1.3 m/s minimizes the net cost of transport. Statistical comparisons made from the regression equations characterizing the non-linear relationship between net metabolic power vs. speed and net cost of transport vs. speed were not statistically different for the time windows taken from 2-4 min, 2-5 min, and 4-7 min (net metabolic power comparisons: *p*_2-4min vs 4-7min_ = 0.431 and *p*_2-5min vs 4-7min_ = 0.543; net cost of transport comparisons: *p*_2-4min vs 4-7min_ = 0.710 and *p*_2-5min vs 4-7min_ = 0.781). Note that the regression curves for the 2-4 min window overlap with the 2-5 min window, therefore, they are indistinguishable from one another. The equations characterizing the polynomial non-linear regression fits in the right column are listed in Table 2.

**Figure 6.**
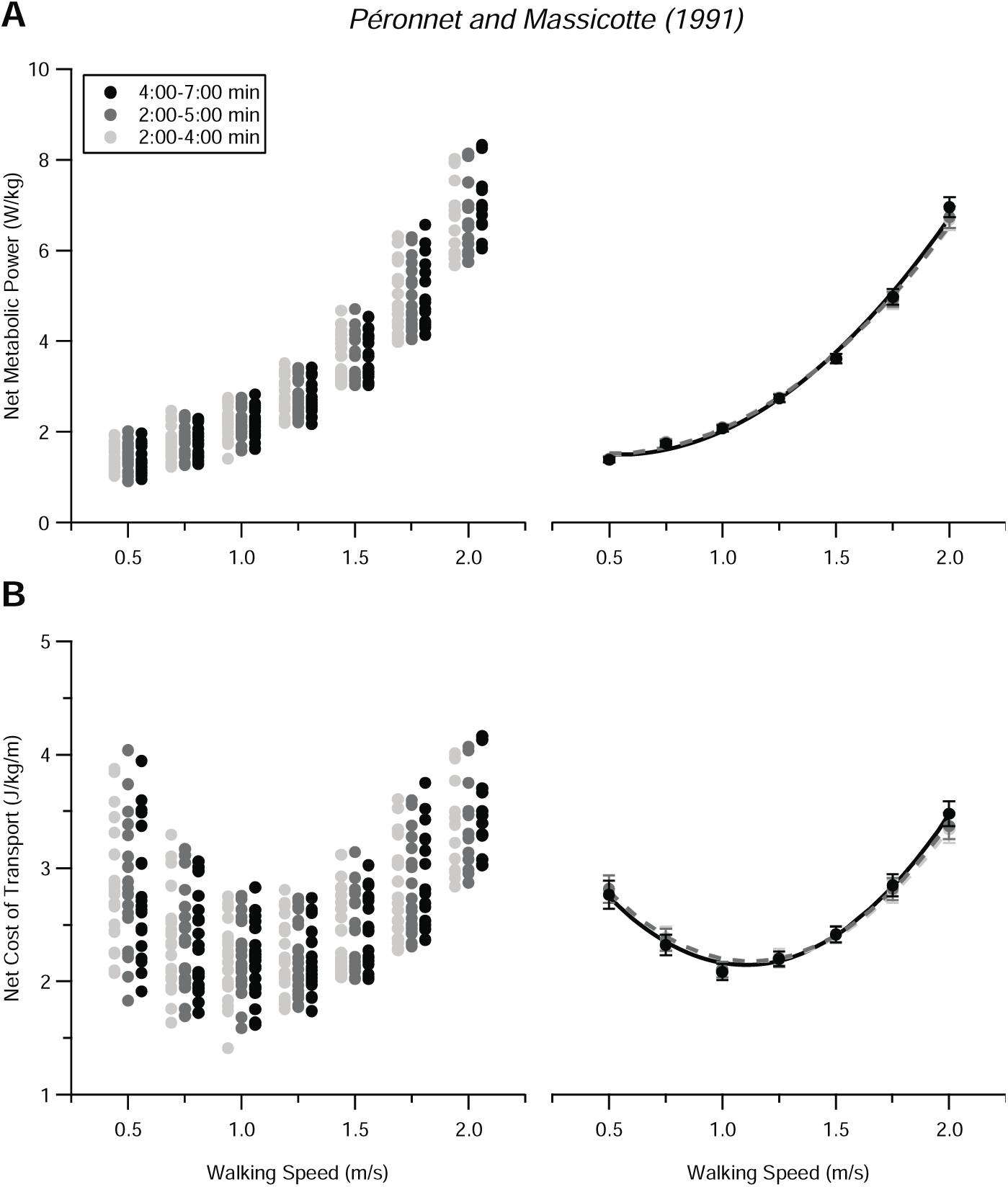
(A) The demand for net metabolic power and (B) net cost of transport changes nonlinearly with walking speed. The left column represents the original data points that are estimated from the Péronnet and Massicotte (1991) equation, and the right column represents the regression curves fit to these data. The data in the right column represent the mean ± sem. Similar to the trends and interpretation in Figure 4 and 5, the demand for net metabolic power increases curvilinearly with speed while the net cost of transport exhibits a U-shaped curve, highlighting the observation that walking at a speed between 1-1.3 m/s minimizes the net cost of transport. Statistical comparisons made from the regression equations characterizing the non-linear relationship between net metabolic power vs. speed and net cost of transport vs. speed were not statistically different for the time windows taken from 2-4 min, 2-5 min, and 4-7 min (net metabolic power comparisons: *p*_2-4min vs 4-7min_ = 0.421 and *p*_2-5min vs 4-7min_ = 0.532; net cost of transport comparisons: *p*_2-4min vs 4-7min_ = 0.699 and *p*_2-5min vs 4-7min_ = 0.774). Note that the regression curves for the 2-4 min window overlap with the 2-5 min window, therefore, they are indistinguishable from one another. The equations characterizing the polynomial non-linear regression fits in the right column are listed in Table 2.

## 4. DISCUSSION

In this paper, we used a slope method as an objective approach to identify when humans achieve steady rates of metabolism while walking across a range of slow to fast speeds (0.5 m/s – 2.0 m/s). We applied this method using a window that quantified the slopes across a 1, 2, and 3-min time period that moved along the time series from beginning to end. In support of our hypothesis, we discovered that at minimum, 4-min of walking at each speed is needed to produce a net COT curve that is not statistically different from a net COT curve obtained from traditional, longer data collection times.

### When do subjects reach a steady rate of metabolism?

Our slope method revealed that across the walking speeds tested here, subjects reached a steady rate of metabolism by 2 minutes, confirming the prediction of Duffy et al. (1996). We came to this conclusion based on our systematic analysis, which provided a clear picture as to how the slope of the net VO_2_ vs time and the slope of net metabolic power vs time changed as each 1-min, 2-min, or 3-min window moved along the time-series data. From Dunnett’s multiple comparison method, one can see that for the 2-min and 3-min intervals (Fig. 3), the *t*-statistic values were highest for the first interval, then decreased steadily until reaching its first minimum value at a time window between 1:30-3:30 min:s and 1:30-4:30 min:s, respectively. Overall, a longer time window of 2-min and 3-min fared much better than a 1-min time window.

### Comparing Net COT Curves for Walking

To keep our comparisons consistent, we calculated the net V□O_2_, net metabolic power and net COT curves derived from a 2-4 min and 2-5 min window. We chose these windows because the regression lines characterizing the change in average slope as a function of speed (Fig. 2) did not differ between a 2-4 min and 4-7 min window and a 2-5 min and 4-7 min window. As illustrated in Fig. 4-6, the net VO_2_, net metabolic power, and net COT curves were not statistically different from the curves produced from the last 3-min of the 7-min trial, our ‘gold’ standard. Follow up comparisons at each speed revealed that pairwise differences between the mean values for net COT (e.g., 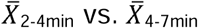 at 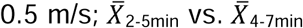 at 0.5 m/s; and so on) were not statistically different, except for the fastest walking speed of 2.0 m/s (Table 1). At this speed, the net COT derived from a 2-4 min and 2-5 min window were slightly larger than our gold standard. We suspect that these differences at 2.0 m/s can be explained by the relatively small sample size of 12 subjects. Nonetheless, including a speed of 2.0 m/s in the net COT curve should be interpreted with caution because this particular speed is where humans prefer to transition from a walk to a run (Farris and Sawicki, 2012; Minetti et al., 1994). This might explain why many of our subjects could not keep up with the treadmill when attempting to walk at 2.0 m/s. If we restrict our walking speeds to a typical range of 0.5 m/s – 1.75 m/s, our analyses suggest that either a 2-4 min and a 2-5 min window will yield average net COT values and curves that are not statistically different from our gold standard.

### Limitations and Future Work

While our slope method was successful for identifying steady rates of metabolism across a range of slow to fast speeds, there are some limitations that warrant further analyses. First, we were unable to determine the effect of sampling rate on identifying periods of steady rates of metabolism. Our data were based on a previous study (Thomas et al., 2021) where our metabolic system was configured to sample average rates of O_2_ consumption and CO_2_ production approximately every 15 seconds. It is possible that applying our slope method on metabolic data that was sampled by breath-by-breath, as done by Schwartz (2007), could have identified shorter time windows, however, breath-by-breath data is inherently noisier. While we show that our results are independent of the typical equations used in the field of locomotion energetics and biomechanics (Brockway, 1987; Péronnet and Massicotte, 1991), a future analysis is needed to address the potential effect of different sampling rates and filtering techniques. Overall, we recommend that any post-processing of these type of data should be kept simple.

In line with the analyses carried out by Schwartz (2007), we also compared our quadratic regression fits and found that the mean square error for the net V□O_2_, net metabolic power, and net COT curves remains small, differing between 6.8% to 12.6% from the ‘gold’ standard (Table 2). This provides evidence that the total time required for data collection can be substantially reduced without sacrificing a decrease in the accuracy of estimating an individual’s net metabolic power and net COT curve. We also recognize that our data are based on a sample of healthy young adults, however, this slope method could be easily applied to any population (e.g., clinical, young children, older adults, etc.) where metabolic measurements are experimentally feasible.

### Conclusion

In summary, we used a slope method as a viable approach for identifying steady rates of metabolism during human walking. Our analyses explored a wide range of slow to fast walking speeds (0.5 m/s-2.0 m/s), revealing that at minimum, 4 min of metabolic data is required to estimate a net COT curve that is not statistically different from a net COT curve derived from traditional data collection times. Our findings suggest that if desirable, shorter trials can be performed, which will help decrease total data collection time. Based on the traditional trial durations that typically range from 6-10 minutes, we estimate a decrease in experimental time of 14-42 minutes across all speeds per subject. For a sample size of 21 subjects, this equates to a decrease of ∼5-15 hours in total data collection time. While it is common for steady rates of metabolism to be confirmed through visual inspection, our slope method provides an objective and simple way to accomplish this experimental goal, which may prove useful for novices and inexperienced researchers who are new to the fields of exercise physiology and locomotion biomechanics.

## Acknowledgements

We thank the University of Houston’s Office of Undergraduate Research who provided support and funding through the Summer Undergraduate Research Fellowship (SURF awarded to B. A.) to complete this project.

## TABLES

## REFERENCES

Alexander, R. M. (1989). Optimization and gaits in the locomotion of vertebrates. Physiological Reviews 69, 1199–1227.

American College of Sports Medicine, Riebe, D., Ehrman, J. K., Liguori, G. and Magal, M. (2018). ACSM’s guidelines for exercise testing and prescription.

Arellano, C. J., McReynolds, O. B. and Thomas, S. A. (2020). A low-cost method for carrying loads during human walking. Journal of Experimental Biology 223,.

Brockway, J. M. (1987). Derivation of formulae used to calculate energy expenditure in man. Hum Nutr Clin Nutr 41, 463–471.

Brooks, G. A., Fahey, T. D. and Baldwin, K. M. (2004). Exercise Physiology: Human Bioenergetics and Its Applications. 4th edition. Boston: McGraw-Hill Education.

Dennis, S. W., Thomas, S. S., Do, K. P., Aiona, M. D. and Wren, T. A. L. (2006). The impact of different normalization schemes on oxygen cost and oxygen consumption data in able-bodied children. Gait & Posture Supplement 2, S286–S288.

Duffy, C. M., Hill, A. E., Cosgrove, A. P., Carry, I. S. and Graham, H. K. (1996). Energy Consumption in Children with Spina Bifida and Cerebral Palsy: A Comparative Study. Developmental Medicine & Child Neurology 38, 238–243.

Farris, D. J. and Sawicki, G. S. (2012). The mechanics and energetics of human walking and running: a joint level perspective. J R Soc Interface 9, 110–118.

Kipp, S., Byrnes, W. C. and Kram, R. (2018). Calculating metabolic energy expenditure across a wide range of exercise intensities: the equation matters. Appl Physiol Nutr Metab 43, 639–642.

Kramer, S. F., Cumming, T., Bernhardt, J. and Johnson, L. (2018). The Energy Cost of Steady State Physical Activity in Acute Stroke. Journal of Stroke and Cerebrovascular Diseases 27, 1047–1054.

Kuo, A. D. and Donelan, J. M. (2010). Dynamic Principles of Gait and Their Clinical Implications. Physical Therapy 90, 157–174.

Maxwell Donelan, J., Kram, R. and Arthur D. K. (2001). Mechanical and metabolic determinants of the preferred step width in human walking. Proceedings of the Royal Society of London. Series B: Biological Sciences 268, 1985–1992.

Minetti, A. E., Ardigò, L. P. and Saibene, F. (1994). The transition between walking and running in humans: metabolic and mechanical aspects at different gradients. Acta Physiol Scand 150, 315–323.

Péronnet, F. and Massicotte, D. (1991). Table of nonprotein respiratory quotient: an update. Can J Sport Sci 16, 23–29.

Plasschaert, F., Jones, K. and Forward, M. (2009). Energy cost of walking: Solving the paradox of steady state in the presence of variable walking speed. Gait & Posture 29, 311–316.

Ralston, H. J. (1958). Energy-speed relation and optimal speed during level walking. Int. Z. Angew. Physiol. Einschl. Arbeitsphysiol. 17, 277–283.

Schwartz, M. H. (2007). Protocol changes can improve the reliability of net oxygen cost data. Gait & Posture 26, 494–500.

Sims, D. T., Onambélé-Pearson, G. L., Burden, A., Payton, C. and Morse, C. I. (2018). The Oxygen Consumption and Metabolic Cost of Walking and Running in Adults With Achondroplasia. Front. Physiol. 9, 410.

Thomas, S. A., Vega, D. and Arellano, C. J. (2021). Do humans exploit the metabolic and mechanical benefits of arm swing across slow to fast walking speeds? J Biomech 115, 1101.1.

Waters, R. L. and Mulroy, S. (1999). The energy expenditure of normal and pathologic gait. Gait & Posture 9, 207–231.

